# Characterization of the growth promotion and defense enhancement of soybean plants driven by seed treatment of multiple soybean-associated beneficial bacteria

**DOI:** 10.1101/2024.01.10.575074

**Authors:** Rosalie B. Calderon, John Christian Ontoy, Inderjit Barpharga, Jong Hyun Ham

## Abstract

We screened soybean-associated bacterial strains from soybean fields in Louisiana, USA, based on various biological activities beneficial for soybean growth and health. Furthermore, we constructed sets of synthetic bacterial community (SBC) containing multiple strains of soybean-associated beneficial bacteria (SABB) having different types of beneficial activities and tested their effects of seed treatment on soybean growth and disease resistance. We found that all three sets of SBC (i.e., Set-1, Set-2, and Set-3) tested promoted soybean growth and yield significantly through seed treatment, showing better performance than the most effective SABB strain *Pseudomonas putida* SABB7 alone and the commercial seed-treating product included for comparison. Our analysis of soybean microbiomes in the root endosphere and rhizosphere based on 16S rDNA sequence profiles revealed that *Bradyrhizobium elkanii*, a symbiotic bacterium of soybean, was enriched in both compartments by seed treatment with Set-2 or Set-m4, which were the best-performing bacterial mixture among the three SABB sets and the most effective subset of Set-2, respectively. In addition, the soybean gene expression profile determined by RNA-seq revealed that seed treatment with Set-2 or Set-m4 made soybean plants grown from the treated-seeds induce a higher level of defense-related genes upon infection by the fungal pathogen *Rhizoctonia solani* compared to those from untreated seeds. These experimental results strongly suggest that the beneficial effects of the bacterial mixtures on plant growth and defense through seed treatment are largely mediated by the change of soybean-associated microbiomes that enriches beneficial components such as *B. elkanii* and the defense-priming effect that induces robust defense responses upon pathogen infection. This study provides a valuable insight into the development of innovative and sustainable management strategies through seed treatment of beneficial microbes in a form of SBC for soybean and further other major crops.

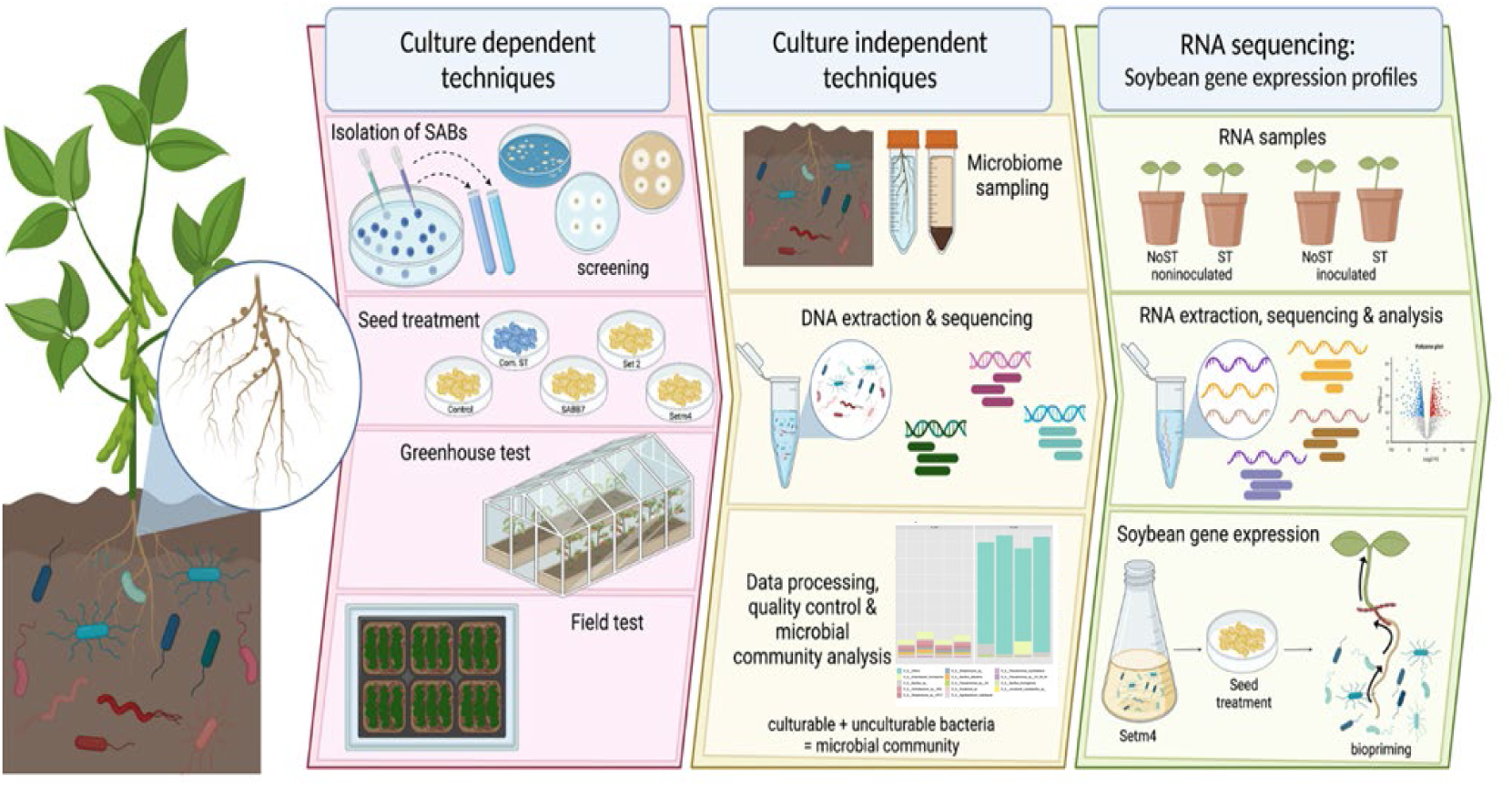

## Introduction

Exploiting beneficial microbes has been considered as a promising strategy to improve crop production with increased sustainability while reducing risks from intensified agricultural practices (1–4). Although beneficial microbes have been extensively explored for growth promotion and protection of crops during the past decades, most of the cases have been dependent on a specific biological activity of a single organism (5–7) (8,9). However, recent studies demonstrated that a group of beneficial microbes could exert a better performance collectively in improvement of crop productivity, which could not be achieved by its individual members alone (2,10–12). In this regard, altering the plant microbiome in its composition or activities for increasing crop performance in nutrient acquisition or tolerance to biotic and abiotic stresses has been proposed as an innovative approach to augment agricultural productivity (1,2).

Soybean (*Glycine max L.*) is the most valuable legume crop in the world, and over one billion metric ton production value of soybeans are produced in the US from nearly 35 million ha of cultivating area, with an estimated annual economic value exceeding $115.8 billion (22). To date, microbiome studies on soybean plants have been mostly about taxonomical characterization of the microbiomes in different plant compartments, such as seed, phyllosphere, root nodules, and rhizosphere (13–15), or those at different soybean growth stages (16). Other topics of soybean microbiome studies include temporal and spatial dynamics of a microbial community affected by crop management practices (14,17,18)(16,19). Although a few studies characterized a natural or synthetic microbial community-bacteria that suppress diseases or increase the efficiency of nutrient uptake through their collective activity (20)(21), to this point, no particular subsets of the soybean-associated microbiome have been linked to overall soybean health and fitness.

The study of plant-associated microorganisms that impact plant functions has recently been focused on systems-level understanding of the collective functionality of microbiome for improving host health and fitness, considering both the plant and the microbiota (1,26–28) (29,30). In soybean, transcriptomic studies using RNA sequencing (RNA-seq) provided a multitude of data sets on plant responses to several biotic and abiotic stresses, such as gene expression of different soybean plant tissues grown in soil inoculated with *Bradyrhizobium japonicum* (31), defense genes against fungal pathogens (*Peronospora manshurica*, soybean downy mildew) and resistance to bacterial pathogens (*Xanthomonas axonopodis pv. glycines,* bacterial leaf pustule) (32–34). Despite these studies, there is limited understanding of how soybean plants express genes differentially in response to seed treatment with beneficial bacteria and infection with soilborne pathogens.

In this study, we observed that seed treatment of bacterial consortia composed of assorted SABB significantly enhanced the disease resistance, growth, and yield of soybean plants in the greenhouse and field conditions. Soybean plants led to significant enhancement of disease resistance and plant vigor in the healthy soybean plants observed in a soybean field with high disease pressure. Further, we investigated the seed treatment effect of a complex group of bacteria with various overlapping beneficial traits in promoting plant growth and immunity and the microbiome structure in soybean roots and the gene expression profile upon pathogen infection in association with the observed disease resistance and growth promotion from the seed treatment of SABB.

## Materials and Methods

### Isolation, screening, and identification of soybean-associated beneficial bacteria (SABB)

The soybean roots and rhizospheric soils from conspicuous healthy plants were collected from the soybean fields at the Doyle Chamber Research Station (30°21’38.6“N 91°10’14.0”W, Ben Hur, Baton Rouge, LA), the Red River Research Station (32°24’56.4“N 93°38’11.0”W, Bossier City, LA), and the H. Rouge Caffey Rice Research Station (30° 14’ 23.0388“ N -92° 20’ 46.0386”W, Rayne, LA) in Louisiana. The soybean-associated bacteria were isolated following the protocol described in (35). The antagonistic abilities of the bacterial isolates were determined by dual-plate confrontation assays as previously described (36). In addition, SABs were screened for growth-promoting activities, including nitrogen fixation, indole-3-acetic acid (IAA) production, phosphate solubilization, siderophore production, and starch hydrolysis following previously described methods (37–40).

Bacterial isolates of SABB that showed strong beneficial activities were identified based on their 16S rDNA sequences. .The DNA was extracted using Qiagen DNA kit (Qiagen, Germantown, MD, USA). The 16S ribosomal DNA was amplified using the bacteria-specific forward primer fD1 (50-AGAGTTTGATCCTGGCTCAG-30) and the reverse primer rP2 (30-ACGGCTACCTTGTTACGACTT-50) (41). The PCR products were sent to MACROGEN, Inc. for sequencing. The DNA sequences were then checked for homology to bacteria-specific genes using the BLAST program of the NCBI-BLAST website (http://blast.ncbi.nlm.nih.gov/Blast.cgi). The phylogenetic tree was constructed using Randomized Axelerated Maximum Likelihood (RAxML) (42).

### *In vitro* co-culture compatibility assay, formulation, and evaluation of bacterial consortia

The best-performing isolates with the most antifungal and growth-promoting activities were subjected to a compatibility test using the cross-streak test described earlier (43). Bacterial strains are considered compatible when streaked colonies grow together, while the zone of inhibition or lysis indicates incompatibility. The bacterial mixture was formulated by mixing compatible strains adjusted to ∼ 1 X 10^9^ CFU/ml (OD_600_=1.0) and combined in a 1:1 ratio (v/v). Subsequently, formulated bacterial consortia were evaluated in the laboratory, greenhouse, and field conditions. Statistical analyses were conducted with JMP Pro Statistics, version 15.1.0 (SAS Institute, Cary, NC).

### Characterization of the soybean microbial community associated with growth promotion

A mesocosm experiment to simulate the soybean microbial community from the field in a more controllable condition was set up in the greenhouse using the field soil from the Doyle Chamber Central Research Station (Baton Rouge, Louisiana), which showed slightly neutral pH and contained moderate organic matter (Table S1). Samplings for characterizing microbiomes in the root endosphere and the rhizosphere were conducted following a previous study (44). The soil samples from the rhizosphere and the root tissue samples for the root endosphere were processed for the downstream DNA extraction procedure using the Qiagen PowerSoil® DNA Isolation Kit (Qiagen, USA). The extracted DNA samples were analyzed for quality and quantity using NanoDrop 1000 (Thermo Scientific, Wilmington, DE, USA). The final DNA samples were sent to the Genomics Research Laboratory of the Biocomplexity Institute of Virginia Tech (Blacksburg, VA, USA) or the Microbiome Services of the University of Minnesota Genomics Center for sequencing, following the 16S Illumina Amplicon Protocol from the Earth Microbiome Project (https://earthmicrobiome.org/protocols-and-standards/16s/). The V4 – V5 region of the 16S rRNA gene was amplified with the 515F/926R primer pair (45,46) and sequenced on the MiSeq platform (Illumina, Inc., San Diego, CA).

The Illumina MiSeq sequencing generated total reads of 3,641,336 for the root endosphere and rhizosphere compartments (Tables S2). The microbial communities were analyzed using the Quantitative Insights into Microbial Ecology (QIIME) pipeline (47). The ‘q2-diversity’ plugin was used to calculate the alpha diversity and beta diversity metrics after samples were rarefied at even sampling depth. The rarefaction curves of the microbiome plateau at the peak indicate a sufficient proportion of diversity represented in the sampling (Fig. S1). Statistical significance corresponding to differences in means and variance among groups was calculated using permutational analysis of variance (PERMANOVA) with 999 iterations. Core microbiome analysis was performed using the core function in R package microbiome (48). Datasets of QIIME output were further visualized using microbiomeAnalyst (49).

### Identification of the SABB strains in the microbiome, differential abundance, prediction of significant and co-occurring species

The SABB strains in the microbiome were identified based on sequence-read similarities. A part of each SBC prepared with pure-cultured SABB was separated right before seed treatment and gone under the same procedure for getting the amplicon sequence profile corresponding to the same V4-V5 region, which was used as a reference sequence indicating the original composition of the SBC and for identification of its SABB components in the microbiome sequences. Based on the sequence read, the ‘species’ taxonomic rank was chosen to identify the SABB strains of the SBC treated, with the highest sequence similarity in the microbiome. Differential abundance of bacterial species was determined using DESeq2 package in R (50). Random Forest (RF) in R package was used to find the most significant taxa in the microbiome (51). The co-occurrence network analysis was performed using the “igraph” package in R (52). A correlation between two amplicon sequence variants (ASVs) was considered statistically significant at > 0.6 of Spearman’s correlation coefficient (r_s_) and < 0.01 of the *p* value (53–55). Multiple testing correction using Benjamini-Hochberg standard false discovery rate correction was used to adjust the *p* values to reduce the chances of obtaining false-positive results (56). Topological properties and statistical analyses were calculated with R using vegan (57) and Hmisc (58) packages. The network was visualized using Gephi platform (http://gephi.github.io/) (59). Bacterial co-occurrence interaction patterns were determined using a python script previously developed (60).

### RNA-seq analysis of soybean seedlings with or without SABB seed treatment in response to Rhizoctonia solani infection

Soybean seeds were surface sterilized before seed treatment with the SBC, Setm4. The seeds were sown in 10cm x 10cm square pots filled with sterilized river silt soil provided by the PMC of LSU AgCenter. A 5 mm-diameter mycelial plug of *R. solani* was placed at the root-shoot junction of a 7-day-old seedling (VC stage) and wrapped with aluminum foil to maintain humidity. Sterile 5-mm-diameter PDA plugs were used for mock inoculation. The four treatments imposed in this study were; mock-inoculated/no seed treatment (M_NoST), mock-inoculated/Setm4 seed treatment (M_ST), R. *solani* inoculated/no seed treatment (T_NoST), and *R. solani* inoculated/Setm4 seed treatment (T_ST). The treatments were arranged in a randomized complete block design (RCBD) in the greenhouse condition. RNA samples were collected at three time points (0-, 3-, and 7-days post inoculation (dpi) with five biological replicates. The collected shoot samples were flash frozen in liquid nitrogen and stored at -80°C until further use.

For total RNA isolation, 100 mg of the powdered (homogenized) soybean tissue was resuspended in 1 ml of TRIzol® Reagent (Ambion® Life Technologies, Grand Island, NY, USA) maintained at 4°C. The resuspended samples were then stored at -80°C until the next RNA extraction steps. The homogenized samples stored in TRIzol® Reagent at -80°C were thawed in ice, and the total RNA of each sample was isolated with the Direct-zol^TM^ RNA MiniPrep Kit (Zymo Research, Irvine, CA, USA) following the manufacturer’s instruction. RNA samples were then treated with DNase I in the Direct-zol^TM^ RNA MiniPrep Kit and finally eluted with 50 μl DNase/RNase-free water.

RNA was normalized to 130 ng/µL in nuclease-free water. RNA integrity was assessed using an Agilent Bioanalyzer 2100 (Agilent Technologies). Libraries were constructed and sequenced using Lexogen Quant-Seq 3’ mRNA-Seq Library Prep Kit (SKU015.96). Library generation was performed using an oligo(dT) primer, and double-stranded cDNA was purified with magnetic beads. Libraries were sequenced on a NextSeq 500 using a high-output single-end, 75 cycles, v2 Kit (Illumina Inc.), resulting in high-quality reads ranging from 11 to 17 million reads per sample (Table S3) by the Genomics Core Facility of Pennington Biomedical Research Center (Baton Rouge, Louisiana). Sequence reads were processed through the Lexogen Bluebee pipeline v 1.8.8, which uses total poly(A) enriched, or rRNA depleted RNA as input (https://www.lexogen.com/corall-total-rna-seq/). The alignment reference was Williams 82 cultivar Assembly 2 (Wm82.a2.v1), and soybean annotation were downloaded from the SoyBase website (https://www.soybase.org/SequenceIntro.php).

The quality of the raw RNA sequences was assessed using FastQC (61). Trimmomatic was used for adapter removal and trimming of bases below the quality threshold level (62). STAR software was used to analyze alignment (63), and featureCounts was used to assign aligned reads to genomic features and generate the reads to count table and summary (64). The DESeq2 V1.24.0 package in R V4.1.2 and Rstudio V1.2.1151 (50) was used for differential expression analysis. Genes with log2FC of >2 (treated/control sample) differences with p-adj 0.1 were labeled as differentially expressed. Significant transcription changes in the RNA-Seq data were determined by applying a False Discovery Rate (FDR) of 0.1. Significant changes were determined after the Benjamini-Hochberg correction for multiple testing (56). Other downstream analyses were performed using the webtool, iDEP.951 Integrated Differential Expression and Pathway (65).

## Results

### Selection of soybean-associated beneficial bacteria (SABB)

A total of 1,740 bacterial isolates were obtained from the root tissues and the rhizosphere of soybean plants, of which 345 strains were screened based on their antifungal and growth-promoting activities. This culture collection included 141 isolates that inhibited *Rhizoctonia solani* in culture (41%), 104 isolates that promoted seedlings growth (30%), 279 isolates that could fix nitrogen (81%), fixed nitrogen, 266 isolates that produced IAA (77%), 41 isolates that produced siderophore (12%) and amylase that can solubilize starch, respectively, and 14 isolates that solubilized phosphate (4%). Among the initially screened 345 bacterial isolates, 31 isolates showing strong and multiple beneficial activities were selected and identified based on their 16S rDNA sequences, which belonged to the genera *Bacillus, Pseudomonas, Streptomyces, Enterobacter, Kosakonia, Leclercia, Ensifer, Rhizobium*, and *Achromobacter*. Nevertheless, none of these SABB exhibited a promising and consistent growth-promoting activity alone through seed treatment in greenhouse or field conditions (data not shown). Thus, we designed a series of SABB mixtures to test if multiple SABB strains would show synergistic or additive effects on the fitness of soybean plants . SABB mixtures were formulated with SABB strains that are compatible among each other based on the co-inoculation assays under the *in vitro* condition (Table S4). We used *Bacillus*, *Pseudomonas,* and *Enterobacter* strains as the base strains because of their strong antagonistic and other beneficial activities. The SABB mixtures tested were composed of five to 23 selected SABB strains (Table S5, Fig. S2), and we use the term ‘synthetic bacterial community (SBC)’ for the SABB mixtures used in this study.

**Figure 1.**
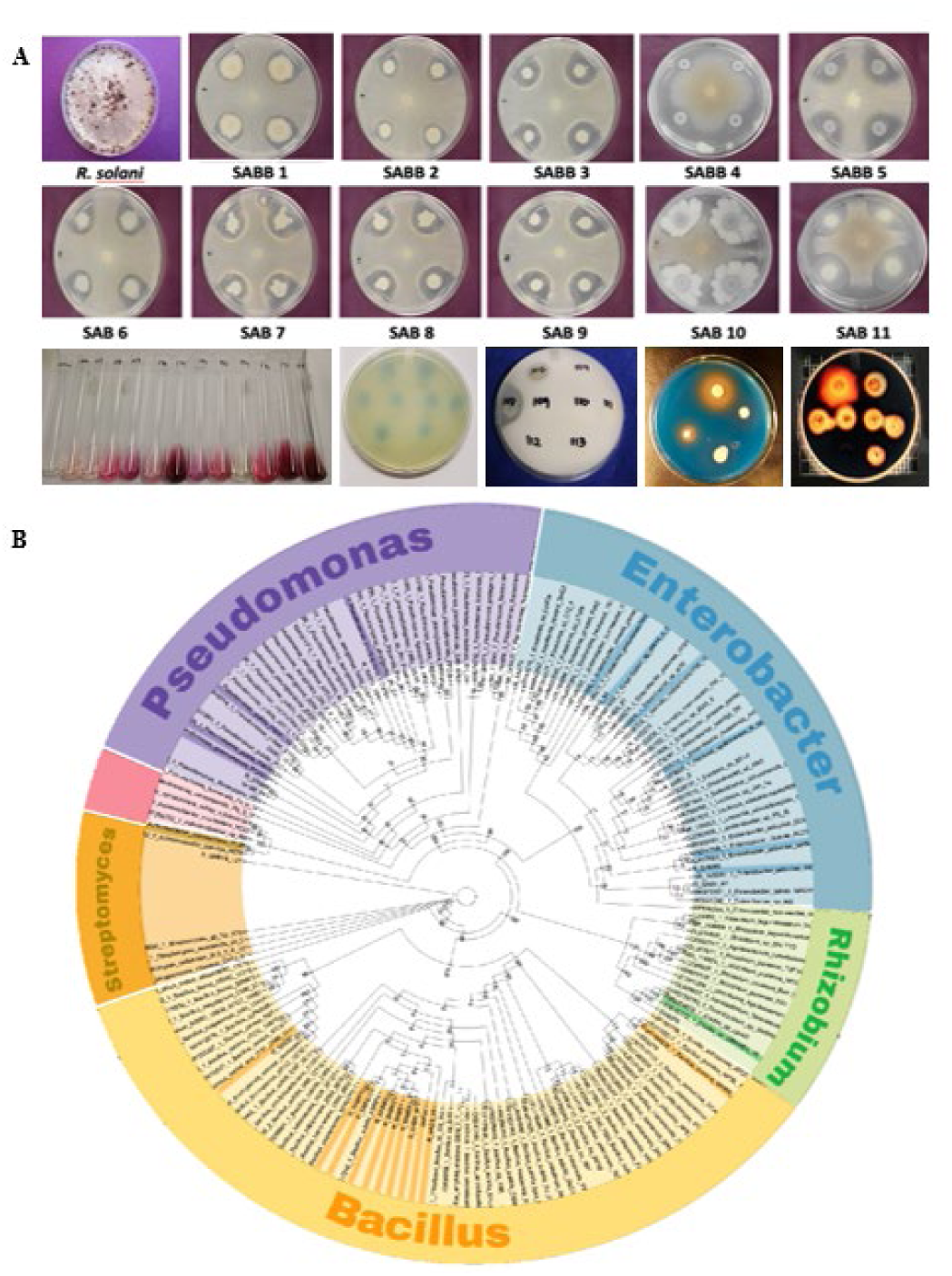
SABB screening A) bioassay against *Rhizoctonia solani,* B) biochemical assays for growth promoting activities: IAA production, nitrogen fixation, phosphate solubilization siderophore production, and starch hydrolysis. (B) Identification in the genus level classification of SABB based on 16S rRNA genes shown in a phylogenetic tree constructed using Randomized Axelerated Maximum Livelihood (RAxML).

### Effect of SABB mixture on seedling root rot, growth and soybean yield

Among the initial seven mixtures evaluated, Set2 and Setm4 treated seedlings showed significantly higher seedling growth based on dry seedling biomass and lesser root rot lesions than the untreated control (Fig. 2A). Both Set2 and Setm4 caused apparently more vigorous growth of soybean seedlings than the commercial fungicide seed treatment, but no significant difference in seedling disease severity was observed in a gnotobiotic assay (Fig. 2A). Further, Set2 and Setm4 treated plants showed significantly higher plant height above ground than the untreated plants in the greenhouse condition using the natural field soil (Fig. 2B *p*=0.00076). Additionally, the nodulation assessed using a WinRHIZO root scanner (Regent Instruments Inc., Canada) indicated Set2- or Setm4-treated plants had a significantly greater number of nodules than the untreated, and Setm4 was more effective than Set2 in promoting nodule formation (Fig. 2B). The number of nodules per plant was significantly correlated with plant growth (Spearman correlation rs= 59%, *p*=0.0103) (Fig. 2B). The soybean yield obtained using a combined harvester in the field showed that plants from the seeds treated with Set2, Setm4, or a commercial seed-treating product (CommST) showed a significantly higher yield than those from untreated seeds (Fig. 2C *p*<0.05).

**Figure 2.**
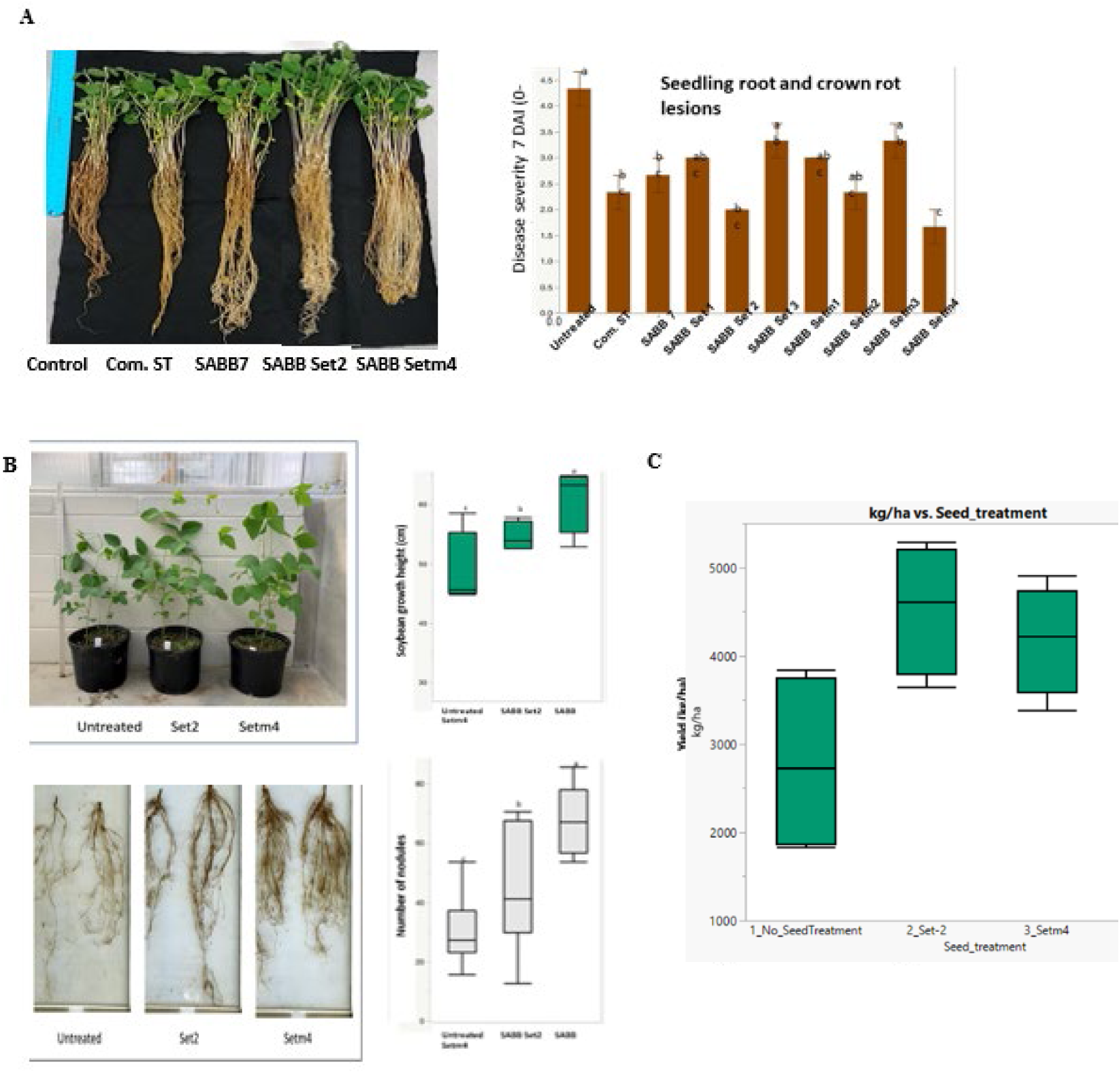
Evaluation of SABB seed treatments. A) seedling root and crown rot assay, B) representative growth and nodulation of SABB Set2 and Setm4 mixture treated plants in comparison with the untreated control and the corresponding boxplots showing the plant height and nodulation of the soybean plants per treatment. C) Soybean yield in a low input field condition. Boxes with a common letter are not significantly different based on LSD tests at 0.05% probability level.

### Characterization of the community structure and core microbiome associated with SBC treated plants

The microbial community of soybean plants grown in a Cancienne silt loam field soil under the greenhouse condition did not vary among the treatments in the level of species richness and Shannon diversity index (*p* >0.05) based on Kruskal–Wallis non-parametric tests (Table S6). There was no distinct variation in the root endosphere microbial structure, nevertheless the rhizosphere structure indicated dissimilarity of the seed treatments from the untreated control based on Bray Curtis dissimilarity as visualized in the PCoA plots (Table S6, Fig. 3, *p*=0.034, R^2^=0.260). A 26% variation in the rhizosphere microbial communities was attributed to the seed treatment effect. Both seed treatments overlap with narrower clusters within the untreated (Fig. 3A). The statistical significance of the clustering pattern in the ordination plot was evaluated using permutational ANOVA (PERMANOVA).

**Figure 3.**
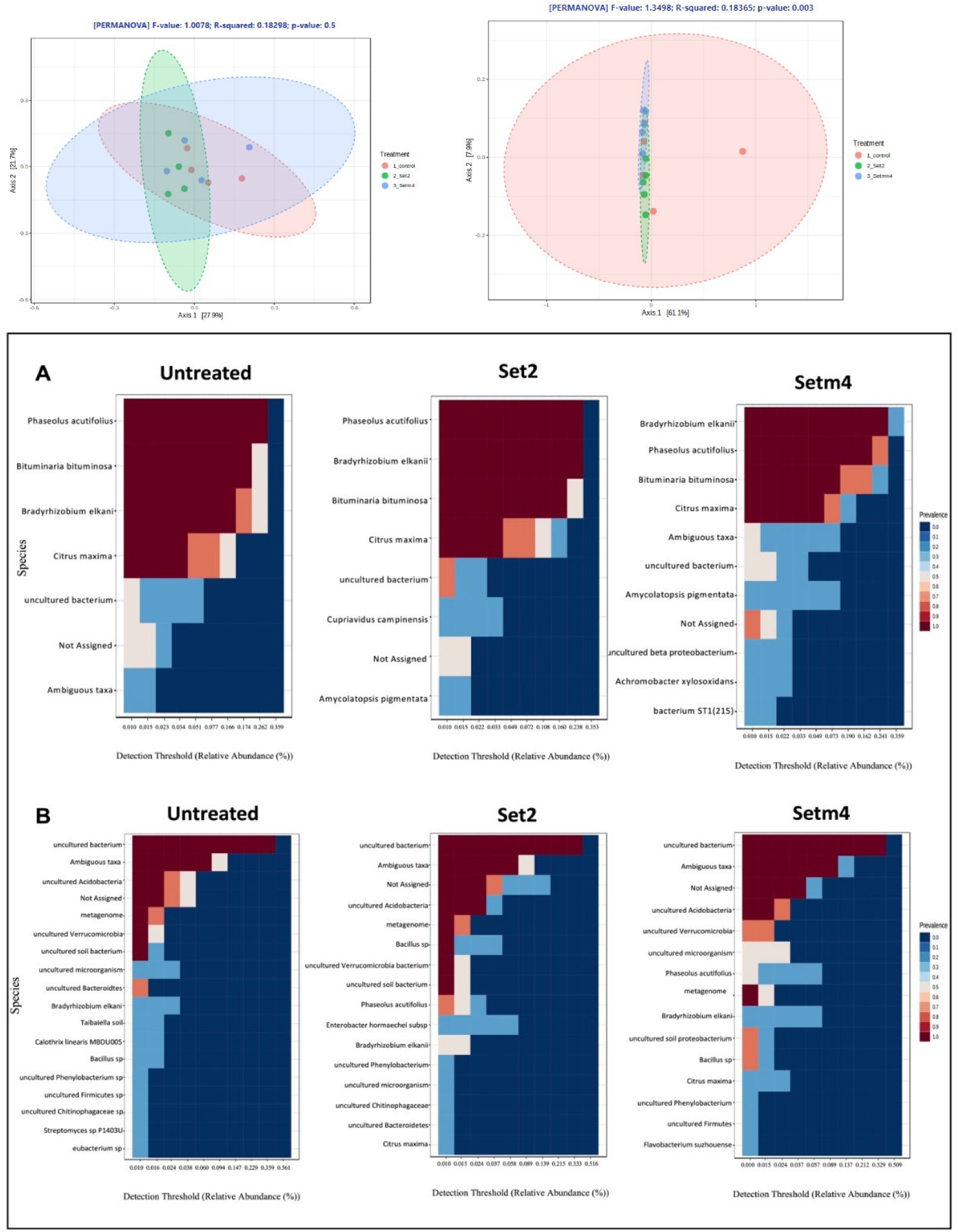
Microbial community structure A) PCoA plots showing the endosphere and rhizosphere compartments, B) Heatmap of core taxa representing the core microbiome at species level of the microbiomes in the root endosphere, and C) the rhizosphere . Y-axis represent the prevalence level of taxa (core features) across the detection threshold (relative abundance) range on x-axis.

The core microbiome in the root endosphere comprised *Bradyrhizobium elkanii*, three species under the phylum *Cyanobacteria* (endophytes in *Phaseolus acutifulus, Bituminaria bituminosa*, and *Citrus maxima*), and uncultured bacterium, which remain unchanged in their composition in the endosphere compartment (Fig. 3B). The important symbiotic soybean bacteria, *Bradyrhizobium elkanii* was more abundant and prevalent in the seed-treated plants with a detection threshold of 0.359 in Setm4 preceded by Set2 treated plants (0.238 detection threshold) than the untreated (0.174 detection threshold) (Fig. 3B). *Achromobacter xylosoxidans,* a strain included in the SBC Setm4, was a distinct core species with 30% prevalence and 0.022 detection threshold in the root endosphere microbiome (Fig. 3B).

Eleven core species across treatments were identified in the rhizosphere microbiome, mostly uncultured bacterium, ambiguous taxa, unassigned species, and metagenome. *Bradyrhizobium elkanii* was prevalent (30 – 50%) in all the treatments like in the root endosphere. It was highly prevalent in Set2 (50% prevalence) and with a greater detection threshold in Setm4 (0.057) than in the untreated (0.024). Similarly, a *Bacillus* sp. and *Enterobacter hormaechei* were more prevalent in Set2 (30-100%) and Setm4 (30 – 70%) with a detection threshold of 0.015 -0.037 (Fig. 3C). The untreated had four core species distinct from the two seed treatments (Fig. 3C).

### Persistence of SABB seed treatment strains, differential abundance, and prediction of significant species in the microbial communities

Based on the sequence read, five strains of the seed-treated SBCs, Setm4 or Set2, were detected in the endosphere microbiomes at the vegetative stage. *Achromobacter xylosoxidans* was detected only in the Setmin4-treated plants, while *Agrobacterium radiobacter* in the Set2-treated plants. The other three strains*, Enterobacter hormaechei, Pseudomonas* sp. D4, and *Bacillus* sp., were detected in both Setm4-and Set2-treated plants (Fig. 4).

In the rhizosphere microbiome, nine strains of the seed-treated SBCs were detected. *Bacillus* sp. and *Pseudomonas* sp. D4 were detected in both setm4-treated and Set2-treated plants. *Bacillus thuringiensis* and *Achromobacter xylosoxidans* were detected in Setm4-treated plants, while *Enterobacter hormaechei, Agrobacter radiobacter, Streptomyces* sp*., Pseudomonas* sp. SV 09 04, and *Ochrobactrum* sp. RS3 (*Rhizobiales*) were in Set2-treated plants (Fig. 4).

**Fig 4.**
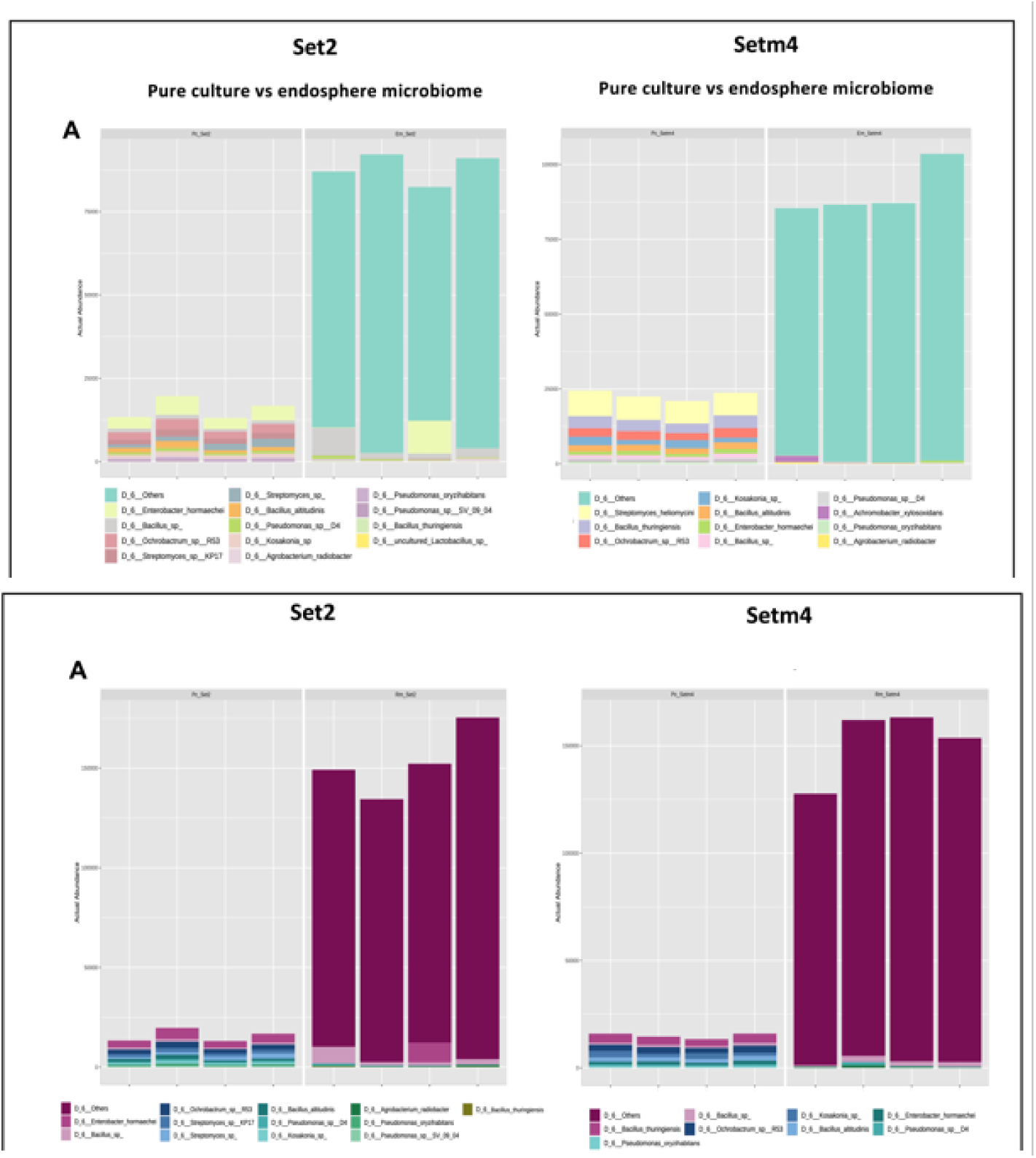
Sequence read comparison of the actual abundance of the SABB pure culture mixtures, Set2 and Setm4 seed treatments in the associated (A) root endosphere and (B) rhizosphere microbial communities at the species level.

Among the differentially abundant species analyzed with DESeq2, the seed-treated SABB *Achromobacter xylosoxidans* was nine folds higher in the root endosphere and six folds higher in the rhizosphere microbiome of Setm4-treated plants than in those of the untreated control plants. *Streptomyces* sp. was three folds greater in the endosphere microbiome of the Set2-treated than in that of the untreated (Table S7).

The strains of the seed-treated SBCs identified as significant species using Random Forest (RF) algorithm were *Achromobacter xylosoxidans* and *Streptomyces* sp. in the root endosphere microbiome. In the rhizosphere, the significant species included A*chromobacter xylosoxidans, Agrobacterium/Rhizobium, Ensifer adhaerans, Rhizobium* sp P26, and *Streptomyces* sp. (Fig. 5).

**Fig 5.**
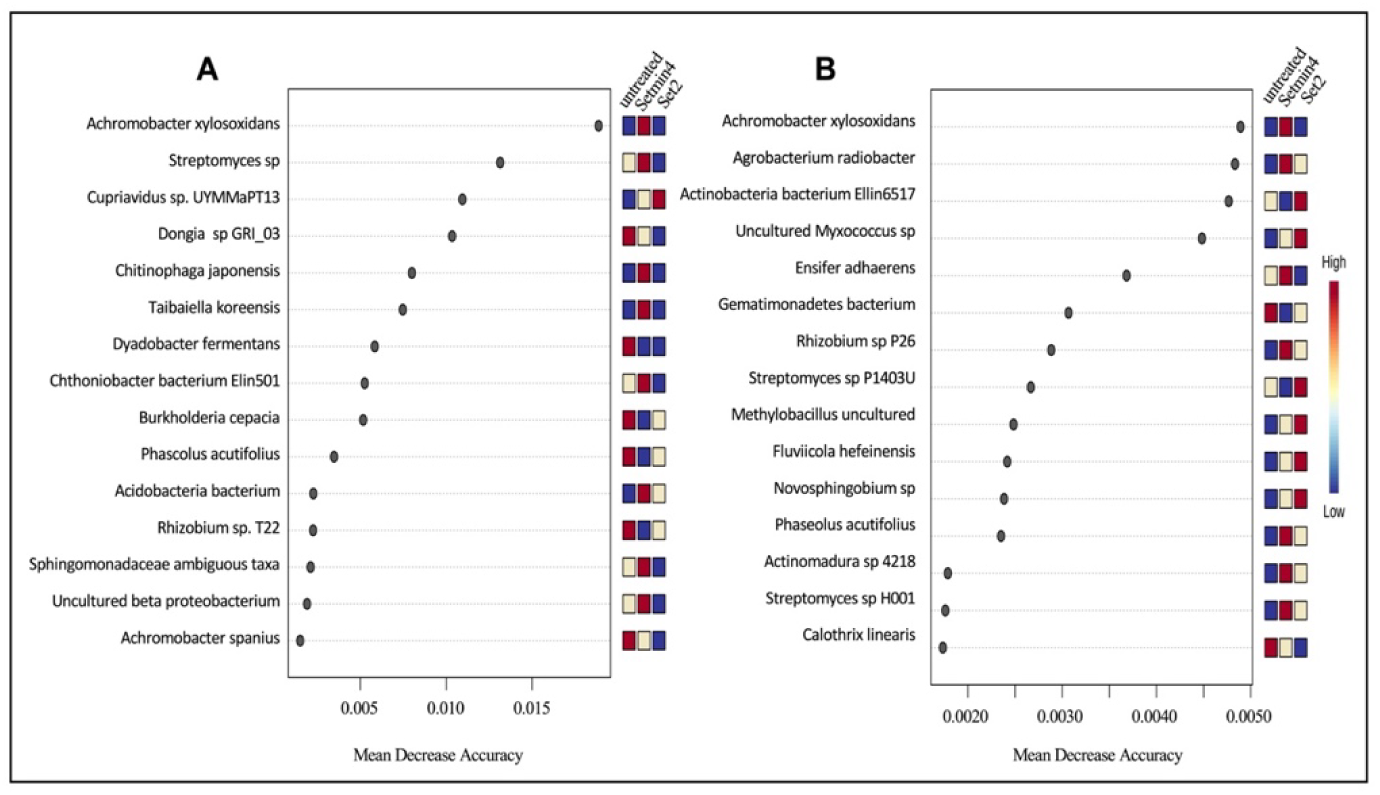
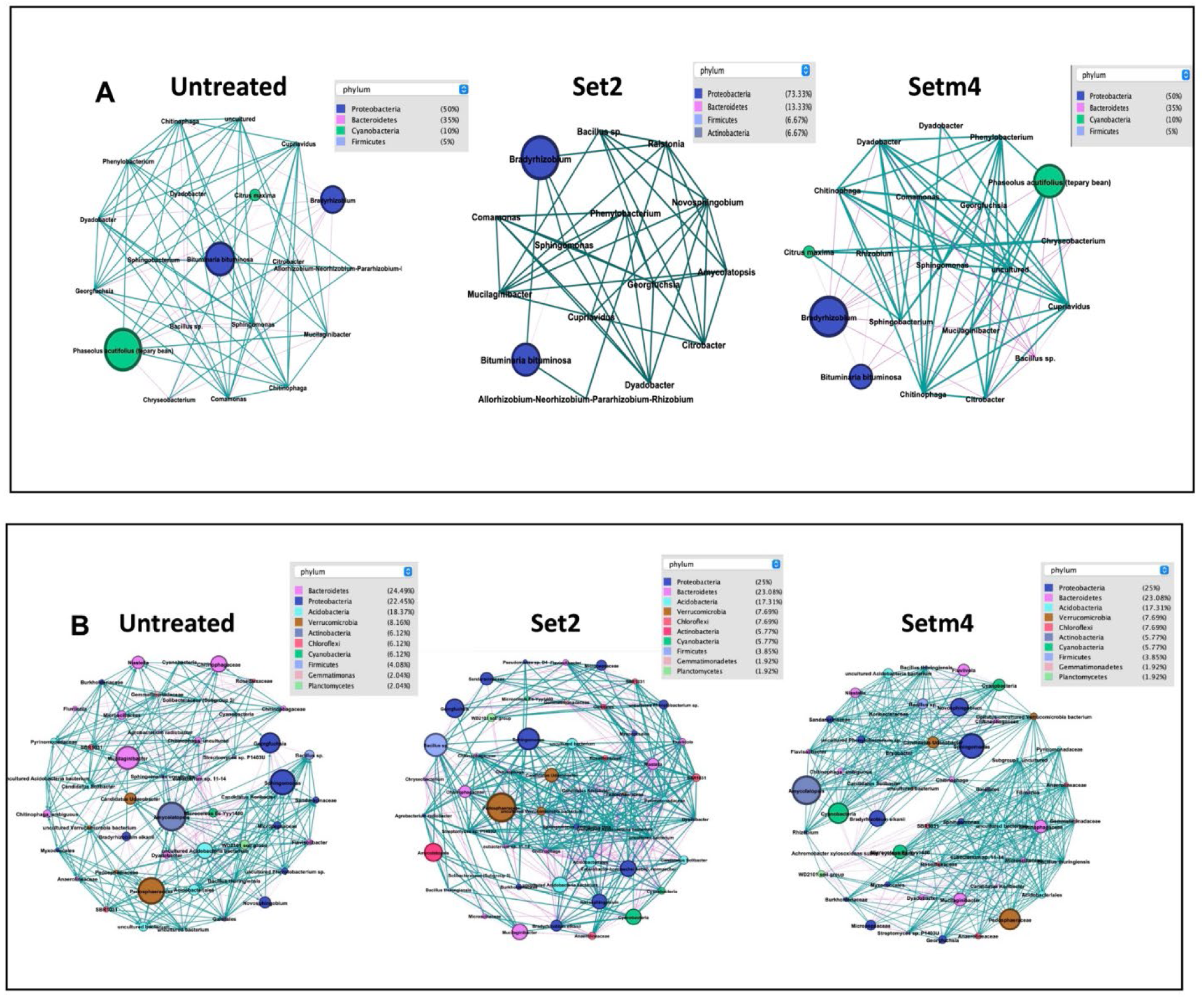
Significant and co-occurring species A) identified by Random Forest in the root endosphere and rhizosphere microbiomes. The features are ranked by the mean decrease accuracy based on permutation and a mini heatmap visualize the patterns of change across the untreated, Setm4 and Set2 seed treatments. B) Microbial interaction networks of dominant taxa at the species level in the endosphere (top 35) and C) rhizosphere microbiome (top 55). The size of the nodes shows taxa abundance, and the different colors indicate the corresponding taxonomic assignment at the phylum level. The edge color represents positive (green) and negative (pink) correlations. The edge thickness indicates the correlation values; only significant interactions are shown (r > 0.6; *p* < 0.01).

Next, we identified key species associated with the seed treatments based on the microbial co-occurrence network using PageRank algorithm. The key species associated with soybean endophytic root microbiome based on relative abundance were *Bradyrhizobium elkaniii*, *Phaseolus acutifolius* (*Cyanobacteria*), and *Bituminaria bituminosa* (*Alphaproteobacteria*) across all the treatments (Fig. 5B). *Bradyyrhizobium elkanii* was more abundant in the seed treated endosphere (Fig. 5B and Table S8).

Similarly, in the rhizosphere microbiome, the seed treatment strains were co-present with known beneficial bacteria belonging to *Rhizobiales*, such as *Bradyrhizobium* sp*, Mesorhizobium* sp., and *Rhizobium* sp. (Fig. 5C and Table S8). The characterized bacterial community structure using a co-occurrence network analysis indicated a nonrandom co-occurrence pattern in both microbiome compartments. The microbial community network showed no significant differences between the seed-treated and the untreated plants in the root endosphere. However, the network in the rhizosphere was more complex and highly dense in the plants treated with Setm4 or Set2 than in the untreated plants (Fig. 5C). This was reflected by the higher number of interacting nodes, edges, average clustering coefficient, average path length, and the greater number of modules or bacterial groups (Figs. 5B, 5C, Table S9).

### Effect of Setm4 seed treatment on the gene expression profile of soybean seedlings inoculated with *Rhizoctonia solani*

Of the 57,147 genes identified using the soybean reference sequence Williams82 a.2.v.1, 39,422 genes were quality filtered and transcriptionally active in the four plant setups: the mock-inoculated or the *Rhizoctonia solani*-inoculated with or without seed treatment with Setm4. Out of the transcriptionally active genes, 7.31% of the expressed transcriptome was differentially expressed, representing 2,882 differentially expressed genes (DEGs) at 0, 3, and 7 days post inoculation (dpi) compared to their controls. The Setm4-treated seedlings had significantly greater biomass than the untreated in both mock-inoculated and inoculated seedlings (P=0.0008) (Fig. 6A). RNA seq analysis revealed upregulation of genes associated with nitrogen metabolism and plant growth hormones in the seed treated (ST) seedlings. Genes involved in nitrate reductase (NR), nitrate/nitrite transporter (Nrt), and carbonic anhydrase (CA 4.2.1.1) were upregulated in the nitrogen metabolism of the plant (Fig. S3). In contrast, genes for ferredoxin-nitrite reductase (nirA 1.7.7.1) and glutamate dehydrogenase (gdha 1.4.1.3) were downregulated (Fig. S3). In addition, the auxin-responsive protein IAA (AUXIAA), auxin-responsive GH3 gene family (GH3), and SAUR family protein for cell enlargement and plant growth were upregulated. Additionally, cytokinin receptor(CRE1), histidine-containing phosphotransfer protein (AHP), and two-component response regulator ARR-B family (B-ARR) in zeatin biosynthesis for cell division and shoot initiation were also upregulated. In contrast, ARF (auxin response factor) and two-component response regulator ARR-A family (A-ARR) were downregulated (Fig. S4).

**Fig. 6.**
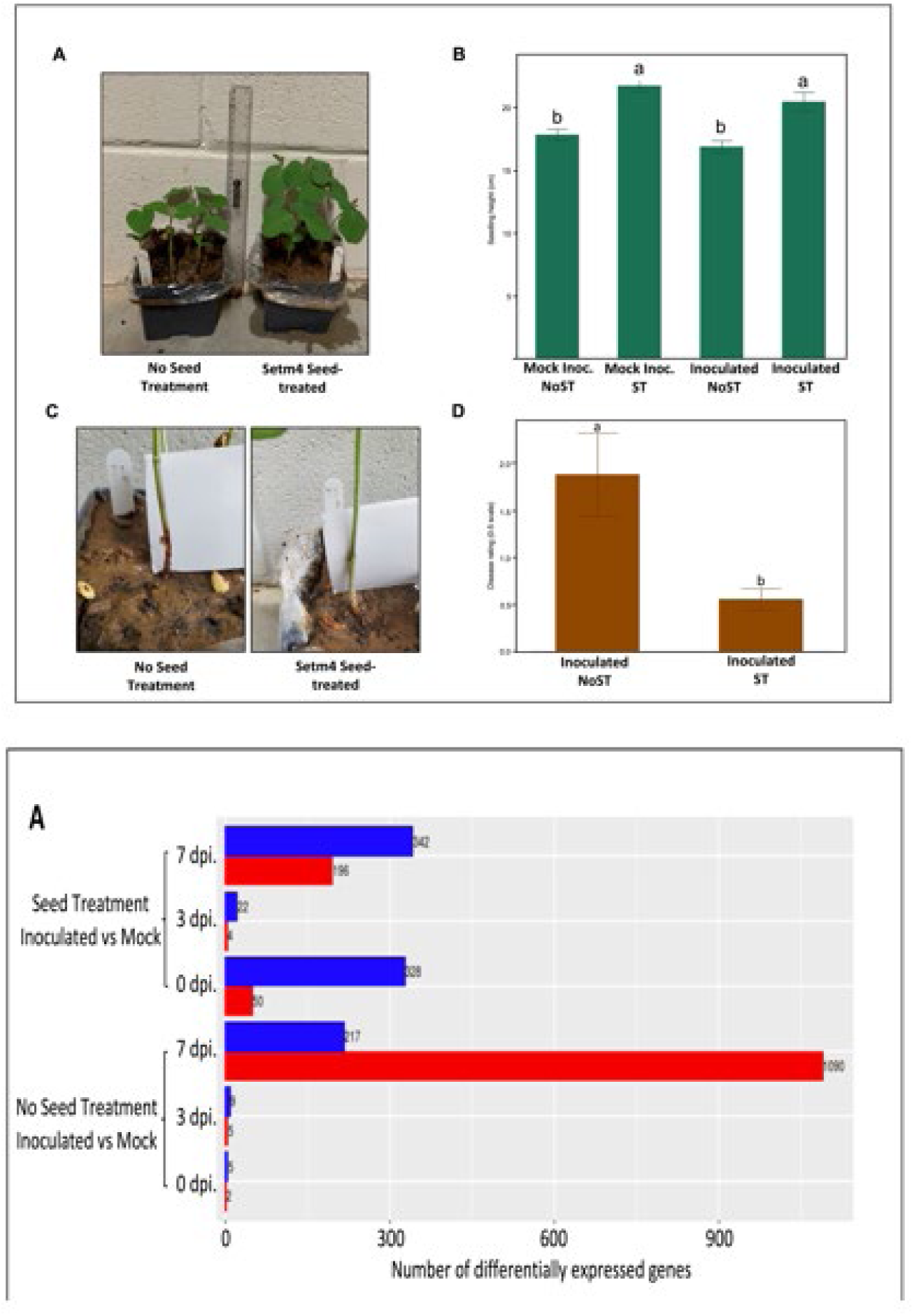
Seedling height and disease severity in response to Setm4 seed treatment and *Rhizoctonia solani* inoculation. (A) Representative soybean seedlings at 12 days after planting. A bar graph showing the height of seedlings among the treatments. Based on oneway ANOVA and the mean bars with common letter are not significantly different at HSD Tukey. Three days old culture of *R. solani* were inoculated by placing 5mm mycelial plugs wrapped with foil around the seedling stem at the VC stage. (B) Representative stem lesions at 7dpi between untreated and seed treated soybeans. Bar graph showing the disease severity ratings in a 0-to-5 scale between the Setm4-treated and the untreated based on Kruskal-Wallis test. (C) Barplot shows the number of differentially expressed genes for each comparison identified by DESeq2. Red bar indicates the up- and blue bar for downregulated DEGs.

The untreated seedlings exhibited reddish necrotic lesions on the stem at or right below the soil line with a greater disease score (P=0.045) than the Setm4-treated at 7 dpi (Fig. 6B). The untreated control with mock inoculation exhibited no disease symptoms (data not shown).

Correspondingly, upregulation of genes related to plant defense included uncharacterized proteins belonging to DEFL family, CRISP, and cytochrome P450. Other genes encoded enzymes, such as xyloglucan endohydrolysis (XEH) and/or endotransglycosylation (XET) glycosyl hydrolase 17 family (hexosyltransferase), tryptophan synthase, sterol desaturase family, SDR, very-long-chain (3R)-3-hydroxy acyl-CoA dehydratase, 3-ketoacyl-CoA synthase, aldehyde dehydrogenase, chitinase class I, peptidase S8 family, aldehyde dehydrogenase, TPP enzymes, xyloglucan endotransglucosylase/hydrolase, peptidase M10A family, lipoxygenase, patatin, and lipolytic acyl hydrolase (LAH). Further upregulated genes included transcription factors, such as AP2-EREBP and NAC (Tables S10, S11). Generally, a lower gene expression level of soybean seedlings was observed with no seed treatment (NoST) in the early time points, but it increased sharply at the later stage of symptom development. In contrast, seedlings with seed treatment (ST) had activated genes early on at 0 dpi, and gradual fluctuations over time (Fig. 6C).

## Discussion

We found that seed treatment of SBC composed of soybean-associated beneficial bacteria (SABB) had a protective effect against soilborne pathogen, *Rhizoctonia solani* and enhances plant growth. We attribute the underlying mechanism of this effect to the persistence of the SABB strains in the microbiome and the biopriming effect from the analyses utilizing 16S rRNA metabarcoding and RNA sequencing data, taking into account both the microbiota and the plant host.

### Plant-associated microbiome harbors beneficial microbes and function together imposing positive effects on plant fitness in natural environment

We obtained a culture collection of more than 300 SABBs with growth-promoting and antagonistic activities against soilborne pathogen from conspicuously healthy soybean plants in the field. Single SABBs had a marginal effect, but SABB consortia had significantly lesser seedling root rot in the lab assay and higher yield in the field conditions. This supports previous studies on the beneficial traits of specific microorganism (6,66–69), but these beneficial traits are an emergent property of the entire microbiome in a natural environment (10,11).

The biocontrol potential of the SBCs could be attributed to the seed treatment strains belonging to *Bacillus* and *Pseudomonas* which are known for producing antibiotics, secondary metabolites, and lytic enzymes reported to act against *R. solani* (70). Other strains comprising the SBCs belong to the genus *Enterobacter*, *Achromobacter, Rhizobium, Ensifer, Streptomyces, Leclercia,* and *Kosakonia*, which are known as plant growth-promoting rhizobacteria capable of N-fixation, phosphate solubilization, IAA and siderophore production, and/or starch hydrolysis. Biological N_2_ fixation could supply N up to 98%, averaging 50 to 60% of the required amount in soybean plants (71). *Bacillus* (72), *Pseudomonas* (73)*, Enterobacter* spp. (74)*, Leclercia* (75), and *Streptomyces* (76) are capable of producing auxin, and most soil bacteria can convert insoluble soil P into plant available forms (77) (78). These SABBs produced siderophores which play a significant role in plant iron nutrition and biocontrol of soil-borne plant diseases (6). They also hydrolyzed starch which indicates their ability to produce amylase and oligo-1,6-glucosidase for nutrient mineralization in the soil (79). These overlapping beneficial traits of the SABBs comprising the SBCs in this study may confer the broader and stronger impacts on plant growth and protection compared to those of single SABB (80)(81)(82).

### Persistence of the seed-treated SABBs and the seed treatment effects of SBCs on microbial communities of soybean plants

At the vegetative stage, persistence of the seed-treated SABB strains in the microbiome was confirmed based on sequence readouts. Along with core species common in all the root endosphere microbiomes, *Achromobacter xylosoxidans* was distinct in the Setm4, which supports that the plant filters and selects the beneficial species that thrive within the plant’s tissue (83). While we observed a higher core species in the untreated than the seed-treated rhizosphere microbiome mirrored in the beta diversity PCoA plot with a significantly narrower cluster of the seed treated than the untreated rhizosphere, which suggests a lesser species composition and restriction of other species that could be attributed to niche occupancy or competition for nutrients (6). This result corroborates with earlier studies of founding taxa at the early plant stage, which have a lasting impact on microbial community assemblies, making them resistant to invasion by latecomers (84). Recent work trying to predict community assembly found that plant-associated microbiomes can be affected by the arrival order, also called the ‘priority effect’, in both assembly and function (85) (86). Here, we observed a similar phenomenon that the seed treatment of SBCs may drive priority effects or lead to a predictable community structure.

The SABB strains of the SBCs were also identified as significant taxa based on the Random Forest. Nevertheless, in our microbiome data, the accuracy and confidence to predict the species classification between the seed treatments and the untreated was 40-60% in the rhizosphere and root endosphere microbiome. The RF model for integrating the abundance of species for phenotypic prediction has been successfully demonstrated in the human microbiome. However, it is more challenging in highly diverse soil than the human gut microbiome, which explains the moderate RF accuracy level (87).

Earlier studies attribute crop productivity as a function of interaction with and among diverse organisms associated with the crop (88–90). In our study, microbial communities associated with soybean growth had a dense microbial network structure which signifies a more complex interaction in the rhizosphere. Other studies reported that microbial network complexity characterizes the high-yield sites compared with the low-yield sites of soybean in the field (91). A non-random co-occurrence pattern inferred in the microbial network interaction indicates that seed treatment strains interact cooperatively with other key species. In this study, the seed-treated strains co-occur with *Bradyrhizobium elkanii*, a symbiotic N fixing bacterium of soybean, which was more abundant in the seed-treated plant compartments. This observation corroborates with other findings that associate the abundance of *Bradyrhizobium* in higher productive soybean field than in areas with lower productivity (92).

### Seed treatment effect of the SBC Setm4 on expression of genes related to plant growth and defense

The gene expression overview revealed a higher level of differential expression in the seed-treated (ST) plants before the challenge inoculation at 0 dpi with more up- and down-regulated genes, suggesting the preconditioning of many genes, which is typical of a ‘primed state’ (93). Specifically, upregulated genes encoding nitrate reductase (NR) is a key enzyme in the assimilation of nitrogen which regulates plant growth (94). Nrt is important in the acquisition of nutrients and their allocation, as well as the ionic balance of nitrate and cellular pH, and is also implicated in the carbon and nitrogen balance (95). This points to the enhancement of N uptake, which provides plants with nutrition favorable for defense. A previous study reported a correlation between low-defense enzymes, such as chitinase, chitosanase, and peroxidase, with low N level (96). Several members of the Nrt family of high affinity nitrate transporter were strongly induced in response to the pathogen (97). The N metabolism genes are strongly affected by pathogen due to defense activation or attempted pathogens manipulation of host metabolism for nutritional purposes. Previous studies suggested that pathogen-resistance-associated NO synthesis influences nutrition homeostasis to favor nitrite diversions either away from infection sites or toward defensive-related metabolism (94,98). Other upregulated defense-related genes in the ST were defensin-like genes encoding DEFL known as antimicrobial polypeptides (99), cysteine-rich secretory protein (CRISP)-related plant pathogenesis proteins of the PR-1 family (100), and chitinase class I against fungal pathogens (Interpro, https://www.ebi.ac.uk/interpro/entry/cdd/CD02877/).

## Conclusion

Plant-associated beneficial bacteria are known to support plant growth and health, but their simultaneous effects on the associated microbial communities and plant gene expression profiles during disease infection remained poorly explored. With 16S rRNA metabarcoding, we show that seed-treated SABB strains persisted in the microbial communities and co-occurred with symbiotic key species, *Bradyrhizobium elkanii*. While RNA sequencing revealed that seed treatment of the SBC Setm4 provided a protective effect on plants by biopriming and inducing a suite of plant defense-related genes upon artificial inoculation of the fungal pathogen *Rhizoctonia solani.* This finding gives a valuable insight into targeted crop management strategies through the application of bacterial inoculants, although their positive effects on host plants in natural settings still remains to be studied more. Taken together, this study not only suggests the seed treatment of SBC as an ideal direction for the practical use of beneficial bacteria for more sustainable cultivation of soybeans but also the scientific basis underlying the positive effects of this crop management practice. Further research is under way to identify core members of each SBC to reduce the number of SABB components without significant loss of beneficial activity for more efficient application of this practice, and to evaluate the effects of SBCs in various field conditions.

## Notes

### Competing Interest Statement

The authors have declared no competing interest.

## References

1. Busby PE, Soman C, Wagner MR, Friesen ML, Kremer J, Bennett A, et al. Research priorities for harnessing plant microbiomes in sustainable agriculture. PLOS Biol [Internet]. 2017 Mar 28 [cited 2021 Sep 25];15(3):e2001793. Available from: https://journals.plos.org/plosbiology/article?id=10.1371/journal.pbio.2001793

2. Chouhan GK, Verma JP, Jaiswal DK, Mukherjee A, Singh S, de Araujo Pereira AP, et al. Phytomicrobiome for promoting sustainable agriculture and food security: Opportunities, challenges, and solutions. Vol. 248, Microbiological Research. Elsevier GmbH; 2021.

3. Singh BK, Trivedi P, Singh S, Macdonald CA, Verma JP. Emerging microbiome technologies for sustainable increase in farm productivity and environmental security. Microbiol Aust. 2018;39(1):17.

4. Singh BK, Trivedi P, Egidi E, Macdonald CA, Delgado-Baquerizo M. Crop microbiome and sustainable agriculture. Nat Rev Microbiol [Internet]. 2020;18(11):601–2. Available from: 10.1038/s41579-020-00446-y

5. De Souza R, Ambrosini A, Passaglia LMP. Plant growth-promoting bacteria as inoculants in agricultural soils. 2015 [cited 2022 Jan 15]; Available from: 10.1590/S1415-475738420150053

6. 6. Lugtenberg B, Kamilova F. Plant-Growth-Promoting Rhizobacteria. 2009; Available from: www.annualreviews.org

7. Qiu Z, Egidi E, Liu H, Kaur S, Singh BK. New frontiers in agriculture productivity: Optimised microbial inoculants and in situ microbiome engineering. 2019 [cited 2022 Jan 13]; Available from: 10.1016/j.biotechadv.2019.03.010

8. Sébastien M, Margarita M-M, Haissam JM. Biological control in the microbiome era: Challenges and opportunities. Biol Control [Internet]. 2015 [cited 2019 Jan 5]; Available from: 10.1016/j.biocontrol.2015.06.003

9. Ray P, Lakshmanan V, Labbé JL, Craven KD. Microbe to Microbiome: A Paradigm Shift in the Application of Microorganisms for Sustainable Agriculture. Front Microbiol. 2020;11(December):1–15.

10. Trivedi P, Leach JE, Tringe SG, Sa T, Singh BK. Plant–microbiome interactions: from community assembly to plant health. Nat Rev Microbiol 2020 1811 [Internet]. 2020 Aug 12 [cited 2021 Jul 6];18(11):607–21. Available from: https://www.nature.com/articles/s41579-020-0412-1

11. Mendes LW, Tsai SM, Navarrete AA, de Hollander M, van Veen JA, Kuramae EE. Soil-Borne Microbiome: Linking Diversity to Function. Microb Ecol. 2015 Jul 28;70(1):255– 65.

12. Finkel OM, Castrillo G, Herrera Paredes S, Salas González I, Dangl JL. Understanding and exploiting plant beneficial microbes. Curr Opin Plant Biol [Internet]. 2017 Aug [cited 2019 Jan 7];38:155–63. Available from: https://linkinghub.elsevier.com/retrieve/pii/S1369526617300158

13. Rascovan N, Carbonetto B, Perrig D, Díaz M, Canciani W, Abalo M, et al. Integrated analysis of root microbiomes of soybean and wheat from agricultural fields. Sci Rep [Internet]. 2016;6(1150):1–12. Available from: 10.1038/srep28084

14. Díaz-Cruz GA, Cassone BJ. Changes in the phyllosphere and rhizosphere microbial communities of soybean in the presence of pathogens. FEMS Microbiol Ecol. 2022;98(3):1–11.

15. Moroenyane I, Mendes & L, Tremblay J, Tripathi B, Yergeau & É. Plant Compartments and Developmental Stages Modulate the Balance between Niche-Based and Neutral Processes in Soybean Microbiome. [cited 2022 Jan 13]; Available from: 10.1007/s00248-021-01688-w

16. Longley R, Noel ZA, Benucci GMN, Chilvers MI, Trail F, Bonito G. Crop Management Impacts the Soybean (Glycine max) Microbiome. Front Microbiol. 2020;11(June).

17. Copeland JK, Yuan L, Layeghifard M, Wang PW, Guttman DS. Seasonal Community Succession of the Phyllosphere Microbiome. Mol Plant-Microbe Interact PMI [Internet]. 2015 [cited 2019 Jan 7];28(3):274–85. Available from: http://dx.

18. Moroenyane I, Tremblay J, Yergeau É. Temporal and spatial interactions modulate the soybean microbiome. FEMS Microbiol Ecol. 2021;97(1):1–12.

19. Calderon RB, Jeong C, Ku HH, Coghill LM, Ju YJ, Kim N, et al. Changes in the microbial community in soybean plots treated with biochar and poultry litter. Agronomy. 2021;11(7):1–26.

20. Mendes R, Kruijt M, De Bruijn I, Dekkers E, Van Der Voort M, Schneider JHM, et al. Deciphering the rhizosphere microbiome for disease-suppressive bacteria. Science (80-). 2011;332(6033):1097–100.

21. Wang C, Li Y, Li M, Zhang K, Ma W, Zheng L, et al. Functional assembly of root-associated microbial consortia improves nutrient efficiency and yield in soybean. J Integr Plant Biol. 2021 Jun 1;63(6):1021–35.

22. JGI Genome Portal. Glycine max v1.0 [Internet]. [cited 2019 Feb 16]. Available from: https://genome.jgi.doe.gov/portal/soybean/soybean.home.html

23. State Agriculture Overview N. USDA/NASS 2017 State Agriculture Overview for North Dakota [Internet]. 2018 [cited 2019 Jan 5]. Available from: https://www.nass.usda.gov/Quick_Stats/Ag_Overview/stateOverview.php?state=LOUISIANA

24. Hartman GL, Rupe JC, Sikora EJ, Domier LL, Davis JA, Steffey KL. Compendium of soybean diseases and pests. Am Phytopath Society; 2015.

25. Lunos AG. Geographic distribution and severity of strobilurin fungicide resistance among Rhizoctonia solani on rice in southwestern Louisiana. Louisiana State University and Agricultural & Mechanical College; 2016.

26. Berendsen RL, Pieterse CMJ, Bakker PAHM. The rhizosphere microbiome and plant health. Trends Plant Sci [Internet]. 2012 [cited 2019 Jan 4];17:478–86. Available from: 10.1016/j.tplants.2012.04.001

27. N Dombrowski KSMASHEKRG-OJWGCPS-L. Root microbiota dynamics of perennial Arabis alpina are dependent on soil residence time but independent of flowering time. ISME J. 2017 Jan 1;11(1):43–55.

28. Trivedi P, Mattupalli C, Eversole K, Leach JE. Enabling sustainable agriculture through understanding and enhancement of microbiomes. Vol. 230, New Phytologist. Blackwell Publishing Ltd; 2021. p. 2129–47.

29. Nyholm L, Koziol A, Marcos S, Botnen AB, Aizpurua O, Gopalakrishnan S, et al. Holo-Omics: Integrated Host-Microbiota Multi-omics for Basic and Applied Biological Research. iScience [Internet]. 2020;23(8):101414. Available from: 10.1016/j.isci.2020.101414

30. Xu L, Pierroz G, Wipf HML, Gao C, Taylor JW, Lemaux PG, et al. Holo-omics for deciphering plant-microbiome interactions. Microbiome. 2021;9(1):1–11.

31. Severin AJ, Woody JL, Bolon Y-T, Joseph B, Diers BW, Farmer AD, et al. RNA-Seq Atlas of Glycine max: a guide to the soybean transcriptome. BMC Plant Biol. 2010;10(1):1–16.

32. Dong H, Tan J, Li M, Yu Y, Jia S, Zhang C, et al. Transcriptome analysis of soybean WRKY TFs in response to Peronospora manshurica infection. Genomics. 2019;111(6):1412–22.

33. Kim KH, Kang YJ, Kim DH, Yoon MY, Moon J-K, Kim MY, et al. RNA-Seq analysis of a soybean near-isogenic line carrying bacterial leaf pustule-resistant and-susceptible alleles. DNA Res. 2011;18(6):483–97.

34. Lanubile A, Muppirala UK, Severin AJ, Marocco A, Munkvold GP. Transcriptome profiling of soybean (Glycine max) roots challenged with pathogenic and non-pathogenic isolates of Fusarium oxysporum. BMC Genomics. 2015;16(1):1–14.

35. Sasser M, Klement Z, Rudolph K, Sands DC. Methods in phytobacteriology. Identif Bact Through Fat Acid Anal Budapest Akad Kaido. 1990;199–204.

36. Shrestha BK, Karki HS, Groth DE, Jungkhun N, Ham JH. Biological control activities of rice-associated Bacillus sp. strains against sheath blight and bacterial panicle blight of rice. PLoS One. 2016;11(1):1–18.

37. Rao S. Soil microorganisms and plant growth. 1977.

38. Gordon SA, Weber RP. Colorimetric estimation of indoleacetic acid. Plant Physiol. 1951;26(1):192.

39. Alexander DB, Zuberer DA. Use of chrome azurol S reagents to evaluate siderophore production by rhizosphere bacteria. Biol Fertil soils. 1991;12(1):39–45.

40. Shruti K, Arun K, Yuvneet R. Potential plant growth-promoting activity of rhizobacteria Pseudomonas sp in Oryza sativa. J Nat Prod Plant Resour. 2013;3(4):38–50.

41. Weisburg WG, Barns SM, Pelletier DA, Lane DJ. 16S ribosomal DNA amplification for phylogenetic study. J Bacteriol. 1991;173(2):697–703.

42. Stamatakis A. RAxML version 8: a tool for phylogenetic analysis and post-analysis of large phylogenies. Bioinformatics. 2014;30(9):1312–3.

43. Santiago CD, Yagi S, Ijima M, Nashimoto T, Sawada M, Ikeda S, et al. Bacterial compatibility in combined inoculations enhances the growth of potato seedlings. Microbes Environ. 2017;32(1):14–23.

44. Edwards J, Johnson C, Santos-Medellín C, Lurie E, Podishetty NK, Bhatnagar S, et al. Structure, variation, and assembly of the root-associated microbiomes of rice. Proc Natl Acad Sci [Internet]. 2015 Feb 24 [cited 2021 Jun 7];112(8):E911–20. Available from: https://www.pnas.org/content/112/8/E911

45. Parada AE, Needham DM, Fuhrman JA. Every base matters: Assessing small subunit rRNA primers for marine microbiomes with mock communities, time series and global field samples. Environ Microbiol. 2016 May 1;18(5):1403–14.

46. Quince C, Lanzen A, Davenport RJ, Turnbaugh PJ. Removing Noise From Pyrosequenced Amplicons. BMC Bioinformatics. 2011 Jan 28;12.

47. Estaki M, Jiang L, Bokulich NA, McDonald D, González A, Kosciolek T, et al. QIIME 2 Enables Comprehensive End-to-End Analysis of Diverse Microbiome Data and Comparative Studies with Publicly Available Data. Curr Protoc Bioinforma. 2020 Jun 1;70(1).

48. Lahti L, Shetty S. Introduction to the microbiome R package. 2018;

49. Chong J, Liu P, Zhou G, Xia J. Using MicrobiomeAnalyst for comprehensive statistical, functional, and meta-analysis of microbiome data. Nat Protoc [Internet]. 2020;15(3):799– 821. Available from: 10.1038/s41596-019-0264-1

50. Love MI, Huber W, Anders S. Moderated estimation of fold change and dispersion for RNA-seq data with DESeq2. Genome Biol [Internet]. 2014 Dec 5 [cited 2020 Nov 25];15(12):550. Available from: http://genomebiology.biomedcentral.com/articles/10.1186/s13059-014-0550-8

51. Liaw A, Wiener M. Classification and regression by randomForest. R news. 2002;2(3):18–22.

52. Csardi G, Nepusz T. The igraph software package for complex network research. InterJournal Complex Syst. 2006;

53. Barberán A, Bates ST, Casamayor EO, Fierer N. Using network analysis to explore co-occurrence patterns in soil microbial communities. ISME J [Internet]. 2012 [cited 2020 Apr 7];6:343–51. Available from: http://www.r-project.org

54. Junker B and FS. Analysis of biological networks. John Wiley & Sons, Inc., Hoboken, New Jersey; 2008.

55. Ju F, Xia Y, Guo F, Wang Z, Zhang T. Taxonomic relatedness shapes bacterial assembly in activated sludge of globally distributed wastewater treatment plants. Environ Microbiol [Internet]. 2014 Aug [cited 2020 Apr 20];16(8):2421–32. Available from: http://doi.wiley.com/10.1111/1462-2920.12355

56. Benjamini Y, Hochberg Y. Controlling the False Discovery Rate: A Practical and Powerful Approach to Multiple Testing. J R Stat Soc Ser B. 1995;57(1):289–300.

57. Ihaka R, Gentleman R. R: A Language for Data Analysis and Graphics. J Comput Graph Stat. 1996;5(3):299–314.

58. Harrell JFE. Hmisc: harrell miscellaneous R package version: 35 – 2 R Foundation for Statistical Computing [Internet]. Vienna, Austria; 2008 [cited 2020 May 12]. Available from: https://github.com/harrelfe/Hmisc

59. 59. Bastian M, Heymann S, Jacomy M. Gephi: An Open Source Software for Exploring and Manipulating Networks Visualization and Exploration of Large Graphs [Internet]. Third International AAAI Conference on Weblogs and Social Media. 2009 Mar [cited 2019 Apr 7]. Available from: www.aaai.org

60. Ju F, Zhang T. Bacterial assembly and temporal dynamics in activated sludge of a full-scale municipal wastewater treatment plant. ISME J. 2015;9(3):683–95.

61. Andrews S, others. FastQC: a quality control tool for high throughput sequence data. 2010. Https://WwwBioinformaticsBabrahamAcUk/Projects/Fastqc/. 2010;7:http://www.bioinformatics.babraham.ac.uk/projects/.

62. Bolger AM, Lohse M, Usadel B. Trimmomatic: a flexible trimmer for Illumina sequence data. Bioinformatics. 2014;30(15):2114–20.

63. Dobin A, Davis CA, Schlesinger F, Drenkow J, Zaleski C, Jha S, et al. STAR: ultrafast universal RNA-seq aligner. Bioinformatics. 2013;29(1):15–21.

64. Liao Y, Smyth GK, Shi W. featureCounts: an efficient general purpose program for assigning sequence reads to genomic features. Bioinformatics. 2014;30(7):923–30.

65. Ge SX, Son EW, Yao R. iDEP: an integrated web application for differential expression and pathway analysis of RNA-Seq data. BMC Bioinformatics. 2018;19(1):1–24.

66. Glick BR. Plant Growth-Promoting Bacteria: Mechanisms and Applications. Scientifica (Cairo). 2012;2012:1–15.

67. Bloemberg G V., Lugtenberg BJJ. Molecular basis of plant growth promotion and biocontrol by rhizobacteria [Internet]. Vol. 4, Current Opinion in Plant Biology. Elsevier Current Trends; 2001 [cited 2019 Oct 16]. p. 343–50. Available from: https://www-sciencedirect-com.libezp.lib.lsu.edu/science/article/pii/S1369526600001837

68. Santoyo G, Moreno-Hagelsieb G, del Carmen Orozco-Mosqueda M, Glick BR. Plant growth-promoting bacterial endophytes. Microbiol Res. 2016;183:92–9.

69. Rashid S, Charles TC, Glick BR. Isolation and characterization of new plant growth-promoting bacterial endophytes. Appl Soil Ecol [Internet]. 2012;61:217–24. Available from: 10.1016/j.apsoil.2011.09.011

70. Cawoy H, Debois D, Franzil L, De Pauw E, Thonart P, Ongena M. Lipopeptides as main ingredients for inhibition of fungal phytopathogens by Bacillus subtilis/amyloliquefaciens. Microb Biotechnol [Internet]. 2015 Mar 1 [cited 2022 Jan 20];8(2):281–95. Available from: https://onlinelibrary.wiley.com/doi/full/10.1111/1751-7915.12238

71. Ciampitti IA, Salvagiotti F. Soybeans and Biological Nitrogen Fixation: A review Seed Yield, Plant N Uptake, and N 2 Fixation Continuing Series: Nutrient Decision Support for Soybean Systems-Part 5. Crops [Internet]. 2018 [cited 2022 Jan 21];102(3):5. Available from: 10.24047/BC10235

72. Shafi J, Tian H, Ji M. Bacillus species as versatile weapons for plant pathogens: a review. Biotechnol Biotechnol Equip [Internet]. 2017;31(3):446–59. Available from: 10.1080/13102818.2017.1286950

73. Rolli E, Marasco R, Vigani G, Ettoumi B, Mapelli F, Deangelis ML, et al. Improved plant resistance to drought is promoted by the root-associated microbiome as a water stress-dependent trait. Environ Microbiol [Internet]. 2015 Feb 1 [cited 2022 Jan 22];17(2):316–31. Available from: https://pubmed.ncbi.nlm.nih.gov/24571749/

74. Kim K, Jang YJ, Lee SM, Oh BT, Chae JC, Lee KJ. Alleviation of salt stress by Enterobacter sp. EJ01 in tomato and Arabidopsis is accompanied by up-regulation of conserved salinity responsive factors in plants. Mol Cells. 2014;37(2):109–17.

75. Graham EB, Knelman JE, Schindlbacher A, Siciliano S, Breulmann M, Yannarell A, et al. Microbes as engines of ecosystem function: When does community structure enhance predictions of ecosystem processes? Front Microbiol. 2016;7(FEB):1–10.

76. Palaniyandi SA, Damodharan K, Yang SH, Suh JW. Streptomyces sp. strain PGPA39 alleviates salt stress and promotes growth of “Micro Tom” tomato plants. J Appl Microbiol. 2014;117(3):766–73.

77. Alori ET, Glick BR, Babalola OO. Microbial phosphorus solubilization and its potential for use in sustainable agriculture. Front Microbiol. 2017;8(JUN):1–8.

78. Anand K, Kumari B, Mallick MA. Phosphate solubilizing microbes: an effective and alternative approach as biofertilizers. J Pharm Pharm Sci. 2016;8(2):37.

79. Pascon RC, Bergamo RF, Spinelli RX, De Souza ED, Assis DM, Juliano L, et al. Amylolytic microorganism from são paulo zoo composting: Isolation, identification, and amylase production. Enzyme Res. 2011;2011(1).

80. Backer R, Rokem JS, Ilangumaran G, Lamont J, Praslickova D, Ricci E, et al. Plant growth-promoting rhizobacteria: Context, mechanisms of action, and roadmap to commercialization of biostimulants for sustainable agriculture. Front Plant Sci. 2018;871:1473.

81. Bashan Y, de-Bashan LE, Prabhu SR, Hernandez JP. Advances in plant growth-promoting bacterial inoculant technology: Formulations and practical perspectives (1998-2013). Plant Soil. 2014;378(1–2):1–33.

82. Saleem M, Hu J, Jousset A. More Than the Sum of Its Parts: Microbiome Biodiversity as a Driver of Plant Growth and Soil Health. Annu Rev Ecol Evol Syst. 2019;50:145–68.

83. Oldroyd GED, Murray JD, Poole PS, Downie JA. The rules of engagement in the legume-rhizobial symbiosis. Annu Rev Genet [Internet]. 2011 [cited 2020 Nov 19];45:119–44. Available from: www.annualreviews.org

84. Carlström CI, Field CM, Bortfeld-Miller M, Müller B, Sunagawa S, Vorholt JA. Synthetic microbiota reveal priority effects and keystone strains in the Arabidopsis phyllosphere. Nat Ecol Evol. 2019;3(10):1445–54.

85. Fitzpatrick CR, Schneider AC. Unique bacterial assembly, composition, and interactions in a parasitic plant and its host. J Exp Bot. 2020;71(6):2198–209.

86. Wei Z, Gu Y, Friman VP, Kowalchuk GA, Xu Y, Shen Q, et al. Initial soil microbiome composition and functioning predetermine future plant health. Sci Adv. 2019;5(9).

87. Blum WEH, Zechmeister-Boltenstern S, Keiblinger KM. Does soil contribute to the human gut microbiome? Microorganisms. 2019;7(9):287.

88. Poudel R, Jumpponen A, Schlatter DC, Paulitz TC, Mcspadden Gardener BB, Kinkel LL, et al. Analytical and Theoretical Plant Pathology Microbiome Networks: A Systems Framework for Identifying Candidate Microbial Assemblages for Disease Management. 2016 [cited 2019 May 10];106(10):1083. Available from: 10.1094/PHYTO-02-16-0058-FI

89. Agler MT, Ruhe J, Kroll S, Morhenn C, Kim ST, Weigel D, et al. Microbial Hub Taxa Link Host and Abiotic Factors to Plant Microbiome Variation. PLoS Biol. 2016 Jan 20;14(1).

90. van der Heijden MGA, Hartmann M. Networking in the Plant Microbiome. PLoS Biol. 2016;14(2):1–9.

91. Bandara AY, Weerasooriya DK, Bell TH, Esker PD. Prospects of alleviating early planting-associated cold susceptibility of soybean using microbes: New insights from microbiome analysis. J Agron Crop Sci. 2021;207(2):171–85.

92. Chang HX, Haudenshield JS, Bowen CR, Hartman GL. Metagenome-wide association study and machine learning prediction of bulk soil microbiome and crop productivity. Front Microbiol. 2017;8(APR):1–11.

93. Mauch-Mani B, Baccelli I, Luna E, Flors V. Defense priming: an adaptive part of induced resistance. Annu Rev Plant Biol. 2017;68:485–512.

94. Fu YF, Zhang ZW, Yuan S. Putative connections between nitrate reductase S-nitrosylation and NO synthesis under pathogen attacks and abiotic stresses. Front Plant Sci. 2018 Apr 11;9.

95. Fan X, Tang Z, Tan Y, Zhang Y, Luo B, Yang M, et al. Overexpression of a pH-sensitive nitrate transporter in rice increases crop yields. Proc Natl Acad Sci. 2016;113(26):7118–23.

96. Dietrich R, Ploss K, Heil M. Constitutive and induced resistance to pathogens in Arabidopsis thaliana depends on nitrogen supply. Plant Cell Environ. 2004;27(7):896– 906.

97. Fagard M, Launay A, Clément G, Courtial J, Dellagi A, Farjad M, et al. Nitrogen metabolism meets phytopathology. J Exp Bot [Internet]. 2014 Oct 1 [cited 2022 Jun 13];65(19):5643–56. Available from: https://academic.oup.com/jxb/article/65/19/5643/2877494

98. Mur LAJ, Simpson C, Kumari A, Gupta AK, Gupta KJ. Moving nitrogen to the centre of plant defence against pathogens. Ann Bot. 2017;119(5):703–9.

99. Tesfaye M, Silverstein KAT, Nallu S, Wang L, Botanga CJ, Gomez SK, et al. Spatio-Temporal Expression Patterns of Arabidopsis thaliana and Medicago truncatula Defensin-Like Genes. PLoS One. 2013;8(3):e58992.

100. Dixon DC, Cutt JR, Klessig DF. Differential targeting of the tobacco PR-1 pathogenesis-related proteins to the extracellular space and vacuoles of crystal idioblasts. EMBO J. 1991;10(6):1317–24.

